# ASGR1 is a candidate receptor for SARS-CoV-2 that promotes infection of liver cells

**DOI:** 10.1101/2022.01.15.476426

**Authors:** Xinyi Yang, Yuqi Zhu, Xiaying Zhao, Jun Liu, Jiangna Xun, Songhua Yuan, Jun Chen, Hanyu Pan, Jinlong Yang, Jing Wang, Zhimin Liang, Xiaoting Shen, Yue Liang, Qinru Lin, Huitong Liang, Min Li, Hongzhou Lu, Huanzhang Zhu

## Abstract

**Backgroud & Aims:** Currently, the COVID-19 pandemic, caused by SARS-CoV-2 infection, represents a serious public health problem worldwide. Although it has been shown that ACE2 serves as the main receptor for SARS-CoV-2 entry into host cells, studies have shown that ACE2 is expressed at extremely low levels in various tissues, especially in some organs where virus particles have been found, such as the heart and liver. Therefore, these organs potentially express additional SARS-CoV-2 receptors that have not yet been discovered.

**Methods & Results:** Here, by a genome-wide CRISPR-Cas9 activation library screening, we found that ASGR1 promoted SARS-CoV-2 infection of 293T cells. In Huh-7 and HepG2 cell lines, simultaneous knock out of *ACE2* and *ASGR1* prevented SARS-CoV-2 pseudovirus infection. In the immortalized THLE-2 hepatocyte cell line and primary liver parenchymal cells, both of which hardly express ACE2, SARS-CoV-2 could successfully establish an infection. After treatment with ASGR1 antibody, the infection rate significantly reduced. This suggests that SARS-CoV-2 infects liver cells mainly through an ASGR1-dependent mechanism. Finally, we also found that the soluble ASGR1 could not only prevent the SARS-CoV-2 pseudovirus, which binds to the ASGR1 receptors, from infecting host liver cells, but also had a protective effect on those expressing ACE2, indicating that administration of soluble ASGR1 protein may represent a new treatment approach.

**Conclusions:** Colletively, these findings indicate that ASGR1 is a candidate receptor for SARS-CoV-2 that promotes infection of liver cells.

**Lay Summary:** We show that ASGR1 is a candidate receptor for SARS-CoV-2 to infect liver cells.

## Introduction

The COVID-19 outbreak, an acute infectious disease caused by severe acute respiratory syndrome coronavirus 2 (SARS-CoV-2), has resulted in a global pandemic, as well as serious public health problems since December 2019 [1]. SARS-CoV-2 is a betacoronavirus with a positive-strand RNA genome [2,3]. It is currently the third coronavirus causing global public health problems; the first two are severe acute respiratory syndrome (SARS) and Middle East respiratory syndrome (MERS) coronaviruses [4,5]. In previous studies, researchers have found that SARS-CoV-2 mainly infects human respiratory cells, especially lung cells [6]. In addition, SARS-CoV-2 could be detected in multiple organs, including the heart, liver, kidney, and intestine [7–11].

Similar to SARS-CoV, researchers found that SARS-CoV-2 enters lung cells mainly through interactions of the viral S protein and the cellular ACE2 receptor [12,13]. Although ACE2 and S proteins display high affinity binding, and cell lines overexpressing ACE2 can be infected by SARS-CoV-2 [13], experimental results obtained by single-cell sequencing showed that many tissues and cells can be infected by SARS-CoV-2, even if the expression level of ACE2 is low. This suggests that there may be cellular receptors for SARS-CoV-2 other than ACE2 [14–16].

The liver is an important organ in the human body.In the process of clinical treatment, researchers have found that some patients with SARS-CoV-2 have visible pathological damage and inflammation in the liver [17,18]. Meanwhile, SARS-CoV-2 was present in patient-derived liver samples upon pathological examination of tissues isolated from corpses [8]. In *in vitro* experiments, multiple liver-derived cell lines could be infected with SARS-CoV-2, but single-cell sequencing results showed that the expression level of ACE2 in liver cells is very low [19]. Therefore, we speculated that liver cells may express a non-ACE2 SARS-CoV-2 receptor.

CRISPR-based genome-wide screening technology has been successfully applied to the screening of multiple receptors involved in viral infection and key host factors [20–22]. In addition, a genome-wide CRISPR knockout library has been used to screen potential host factors important in SARS-CoV-2 replication [23,24]. This may be intrinsic to the selected cell model not expressing the SARS-CoV-2 non-ACE2 receptor; hence, screening of the CRISPR knockout library could not be employed to get a positive result. Therefore, to overcome the above-mentioned obstacles, we used CRISPR-activated libraries for undifferentiated SARS-CoV-2 non-ACE2 receptor screening.

## Materials and Method

### Antibody and reagents

The following antibodies were used throughout this study: from ABclonal(Wuhan, China), anti-FLAG (AE063), anti-HA (AE036), anti-ASGR1 (A13279), anti- β -Actin (AC026). From Abcam (Cambridge, UK), anti-ASGR1 (ab254262). From proteintech (Wuhan, China), anti-ACE2 (21115-1-AP). Anti-Flag Magnetic Beads (HY-K0207) was purchased from MCE. 2 × Taq Master Mix (P112), High fidelity PCR enzyme- 2 × Phanta Max Master Mix (P515) were purchased from Vazyme (Nanjing, China). PMD18-T (6011) was purchased from Takara (Beijing, China). Cell Genome Extraction Kit (DP304), Plasmid Extraction Kit (DP103, DP108, DP117) were purchased from Tiangen (Beijing, China). Gel Extraction Kit (CW2302) was purchased from CWBIO (Nanjing, China). Luciferase detection kit (E6110) was purchased from Promega (Madison, USA). Cell Counting Kit (CCK-8) and TUNEL Apoptosis Detection Kit (FITC) TUNEL kit were purchased from Yeasen (Shanghai, China).

### Cell culture

293T, HeLa, HepG-2, Huh-7, H1299, A549, BEAS-2B, cells were were cultured in DMEM (Gibco, C11995500BT) with 10% fetal bovine serum (FBS) (ExCell Bio, FSP500) and 1% penicillin/streptomycin (P/S) (Gibco, 15140-122) in a 37 ° C incubator containing 5% CO_2_. CaCo-2 cells were cultured in 1640 (Gibco, C11875500BT) and supplemented with 10% FBS (ExCell Bio, FSP500), and 1%P/S in a 37 °C incubator containing 5% CO_2_. THLE-2 and primary hepatocytes cells were cultured BEGM (Lonza, CC-3170) in with 5 ng/mL EGF, 70 ng/mL Phosphoethanolamine and 10% FBS in a 37 °C incubator containing 5% CO_2_

### Vector construction and cas9-mediated gene knockout and cDNA overexpression

Individual sgRNA constructs targeting ASGR1 or ACE2 was cloned into lentiCRISPR v2.0 (addgene 52961). For cDNA expression vectors, a linearized lentiviral backbone was generated from PCDH (Youbio, Hunan, China). Protein-coding sequences were gift from MiaoLing Plasmid Sharing Platform. All the constructed plasmids were confirmed by restriction enzyme digestion and DNA sequencing. Huh-7 or 293T cells were infected with lentivirus at an MOI of 1 and then selected using 2 μg/ml puromycin for 14 days for knockout or overexpression. The knockout efficiency was analyzed using Sanger DNA sequencing. Knockout efficiency was detected by western blot (WB) analysis.

### Construction and production of pseudoviruses

The spike of SARS-CoV-2 and that of all variant plasmids were synthesized by the GeneScript Company. All S proteins were optimized with 18 amino acids removed and an HA tag attached. The core plasmid, pLenti.GFP.NLuc-Puromycin, that expresses GFP, luciferase and puromycin at the same time was maintained in our laboratory.

For production of pseudoviruses, HEK293T/17 cells were seeded into 10cm dishes one day before transfection. When the cells reached 80% confluence, plasmid and PEI were added into the Opti-MDM (Gibco), mixed evenly, and left standing for 20min. pLenti.GFP.NLuc, psPAX2 and the S protein vector were co-transfected into HEK293T/17 cells to produce the pseudoviruses. After incubation at 37°C and 5% CO_2_ for 24h, the culture medium was changed with DMEM with 2% FBS and 1% P/S. The supernatants containing SARS-CoV-2 pseudotyped viruses were harvested at 48h and 72h after transfection and filtered by 0.45 μm pore size. The filtrates were centrifuged at 25000rpm and 4°C for 2h. The supernatants were discarded, and the pseudovirus stocks were dissolved in DMEM for storage at −80°C [25–27].

### Visualization of GFP and flow cytometry assay

The cells were collected and washed with phosphate buffered saline (PBS). They were kept in PBS before analysis on a BD LSRII flow cytometer for EGFP expression. FlowJo software (FlowJo LLC, Ashland, OR) was used to perform the flow cytometry analysis.

### Luciferase reporter assay

The cells were collected and washed with phosphate buffered saline (PBS). Cells were harvested at 72 hours post-infection, and the lysate was assayed for luciferase activity. Triplicate cultures were measured for each experiment.

### Western blot

A total of 1 × 10^6^ cells were preseeded in 10-cm dish and cultured 24 h. Then cells were harvested, lysed, and subject to SDS PAGE, then transferred on N.C membrane, followed by the incubation with indicated primary antibody. Membranes were visualized using the Immun-Star WesternC Chemiluminescence Kit (Bio-Rad) and images were captured using a ChemiDoc XRS+ System and processed using ImageLab software (Bio-Rad).

### Protein purification

The ASGR1 protein was produced in HEK 293T cells. After the plasmid of S protein was transfected for 48 h, the cells were collected and lysed. Anti-Flag protein A/G magnetic beads were then used for immunoprecipitation and quantified by BCA kit.

### Cell proliferation by CCK-8 assay

Five thousand cells were added to each well of 96-well plates. To measure the OD value, 10% CCK8 solution was added to fresh culture medium and incubated at 37°C for 1 hour. The OD 450 nanometer value was measured.

### Apoptosis detected by Tunel staining

A total of 1 × 10^6^ cells were collected in a 1.5 ml tube and centrifuged at 300 g for 5 min. The cells were washed twice with 500 μ L PBS. Cells were treated with TUNEL-FITC apoptosis detection kit according to the manufacturer's instructions. The proportion of FITC-positive cells in the cells was analyzed by flow cytometer. FlowJo software (FlowJo LLC, Ashland, OR) was used to perform the flow cytometry analysis.

### Statistical analysis

Data are representative of three independent experiments, and error bars represent standard errors (SD). Paired samples t-tests were performed with use of SPSS version 13.0 (SPSS Inc., Chicago), and statistical significance was indicated at *P < 0.05, **P < 0.01 or ***P < 0.001.

## Results

### Construction and identification of SARS-CoV-2 pseudovirus based on a lentiviral system

Since authentic SARS-CoV-2 must be operated in a P3 laboratory and lacks a standard reporter gene, it was not conducive to our screening approach. We first constructed an expression plasmid encoding the viral S protein, in which we replaced 18 amino acids at the C-terminus by an HA tag (Fig. 1A) [28]. To verify the expression of the S protein, we analyzed total protein levels of 293T cells transfected with S protein- and vesicular stomatitis virus G (VSVG) protein-encoding plasmids by western blotting (Fig. 1B). Next, we used 293T cells to assemble VSVG and S proteins into capsid proteins to package pseudoviruses. The collected pseudoviruses were analyzed using antibodies against the HA-tag and serum obtained from people immunized with inactivated SARS-CoV-2 vaccines (Fig. 1C-1D), We further observed the structure of the pseudovirus using electron microscopy (Fig. 1E). These results confirm that we have successfully constructed virus particles containing S protein. Finally, we used pseudoviruses loaded with green fluorescent protein (GFP) and luciferase reporter genes packaged by the viral S protein to infect different cell lines *in vitro*. The results showed that the packaged pseudoviruses not only infected 293T cell lines overexpressing ACE2, but also Huh-7, HepG-2, H1299, and CaCo-2 cells. However, the SARS-CoV-2 pseudovirus cannot infect cell lines such as HeLa. (Fig. 1F-1G). This was consistent with infection experiments reported by other research groups [29], suggesting that we successfully constructed a SARS-CoV-2 pseudovirus packaged by the S protein based on a lentiviral system.

**Figure 1.**
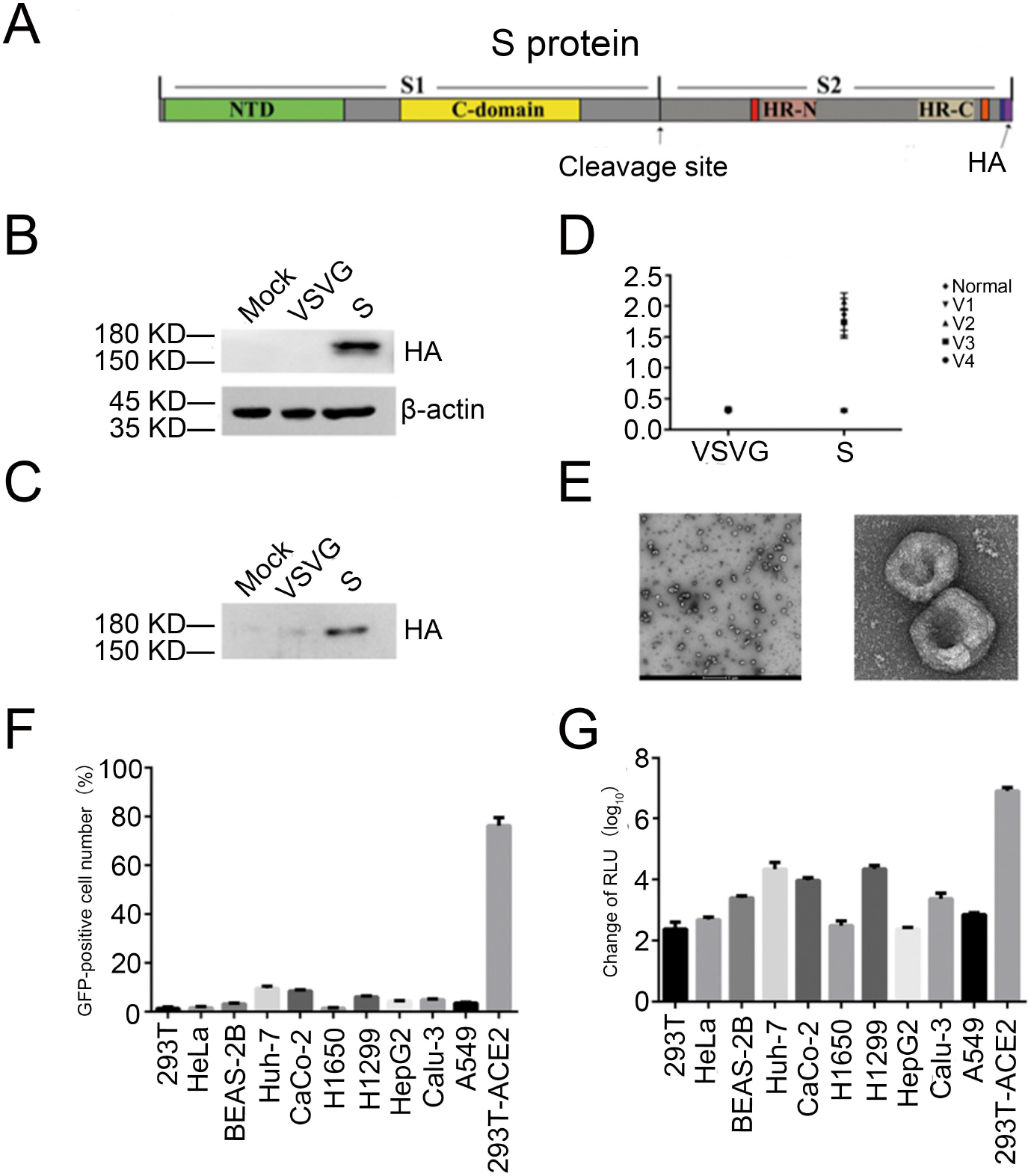
Construction and verification of SARS-CoV-2 pseudovirus based on lentivirus system. (A) Schematic diagram of the S protein vector for packaging SARS-CoV-2 pseudovirus. (B) The expression of S protein were measured by Western blot in 293T cells transfected with the plasmid encoding S protein or VSVG protein. (C) The expression of S protein were measured by Western blot in virus supernatant of S protein or VSVG packaged. (D) The pseudovirus packaged by S protein has neutralizing activity. The sera of the vaccinators are incubated with the pseudovirus, and the neutralizing activity is tested by ELISA. (E) Morphological detection of pseudovirus packaged by S protein. After the prepared pseudovirus was negatively stained, it was observed using a projection electron microscope. (F, G) S-protein packaged pseudoviruses infect different cell lines. Incubate the pseudovirus with 293T, HeLa, Huh-7, BEAS-2B, Caco-2, H1650, Calu-3, 5-8F, HepG2, A 549 and 293T-ACE2 cells for 12 hours, replacing the fresh culture and 72 hours later, the expression levels of GFP (F) and luciferase (G) were detected by cell flow cytometer and microplate. Each data represented the mean ± SD of three independent experiments (n=3) and were analyzed with T-test.

### Genome-wide CRISPRa screening identifies new potential receptor for SARS-CoV-2

To identify viral entry factors for SARS-CoV-2, we established SAM library screening [30] in HeLa cells, which has been proven by us and other researchers that it cannot be infected with the SARS-CoV-2 pseudovirus [12] (Fig. 1F-1G). This SAM library contains over 70920 gRNAs for 23430 human genes [30]. We prepared MPH lentivirus to infect HeLa cells with a multiplicity of infection (MOI) of 10, after which the cells were subjected to hygromycin (200 μg/ml) selection for 14 days (Fig. 2A). We extracted the genome of the selected cell line and identified it by PCR using primers specifically targeting the MPH plasmid region. We found that infected and drug-screened cell lines had integrated the p65 gene sequence in MPH management (Fig. 2B). We also prepared a SAM library lentivirus to infect HeLa-MPH cells with an MOI of 0.2, with the aim of single cells being infected by only one virus particle. Then, the cells were selected with blasticidin, and Cas9 expression was analyzed by western blotting with anti-Cas9 antibody (Fig. 2A and 2C).

**Figure 2.**
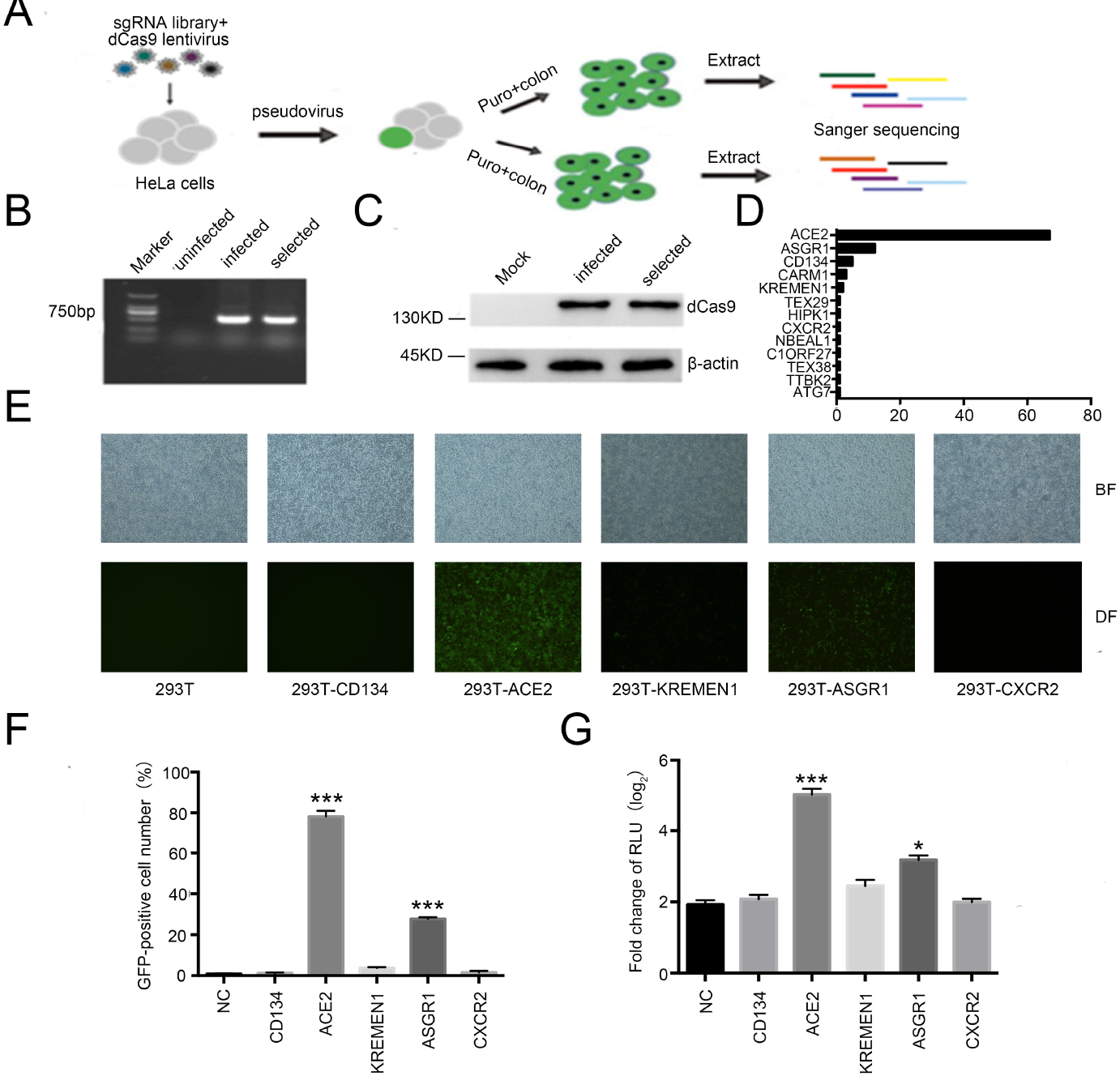
A pooled, genome-wide CRISPR screening for candidate receptor involved in SARS-CoV-2 entry. (A) The outline of genome-wide activation CRISPR screen strategy. The HeLa cells were infected with lentiviral containing MS2-p65-HSF1 proteins treated with Hygromycin B for fourteen days. Then, the HeLa cells expressing MS2-p65-HSF1 infected with lentiviral dCas9 and sgRNA library that target 20234 human genes. Fourteen days post screening with blascicidin, genomic DNA was extracted from monoclonal cells after puromycin selected. The candidate genes were identified by the Sanger sequencing. (B) Validation of MS2-p65-HSF1 in uninfected, infected and HeLa cells with PCR after infection. (C) After infected with the HeLa cells expressing MS2-p65-HSF1 with SAM library, western blot of cell lysate with anti-Cas9 antibody. (D) Statistical analysis of candidate sgRNA. The sgRNA regions of all monoclonal cells were subjected to Sanger sequencing, and the candidate genes corresponding to sgRNA were ranked according to abundance. (E-G) ASGR1 facilitates SARS-CoV-2 virus pseudotype infection as potently as ACE2. HEK293T cells were transfected with Flag-tagged CD134, ACE2, KREMEN1, ASGR1, CXCR2 or empty, then infected with the GFP-labeled and luciferase SARS-CoV-2 virus pesudotype, and visualized by microcopy (E), the expression levels of GFP by cell flow cytometer (F), luciferase were detected by microplate (G) at 72 h post infection. Each data represented the mean ± SD of three independent experiments (n=3) and were analyzed with T-test. *, p < 0.05;***, p > 0.001.

After preparing the cell library, we prepared a pseudovirus containing a GFP-luciferase-puromycin construct packaged with S protein to infect the cells from our library at an MOI of 10. Next, puromycin was added to select cells that had been successfully infected (Fig. 2A). After the screening process, we found that most of the cells had died, and the surviving cells grew into single clones. Therefore, we expanded these surviving monoclonal cell populations, extracted the cellular genome, and sequenced the sgRNA sequences present in these monoclonal cells by Sanger sequencing (Fig. 2D). Through statistical analysis, we obtained a total of 13 sgRNAs targeting different genes, including *ACE2*. In addition to *ACE2*, we found the genes encoding CD134, KREMEN1, ASGR1, and CXCR24 membrane proteins (Fig. 2D). Considering that SARS-CoV-2 enters cells mainly through interaction of the S protein and membrane proteins, we focused on the above-mentioned membrane proteins for subsequent experiments.

### Validation of candidate genes encoding factors for SARS-CoV-2 entry into cell

To test whether these candidate genes were relevant to SARS-CoV-2 entry into the cells, we infected 293T cells with lentiviruses carrying different candidate genes, followed by clonal selection with puromycin (2 μg/ml) for 14 days. Next, we used S protein-packaged pseudovirus containing GFP and luciferase to infect the aforementioned cell lines. Using fluorescence microscopy, we found that in addition to 293T-ACE2 cells, 293T-ASGR1 cells were susceptible to infection with SARS-CoV-2 pseudovirus (Fig. 2E). We further confirmed these results through flow cytometry and luciferase reporter gene analysis (Fig. 2F-2G). These results suggest that ASGR1 may be a novel SARS-CoV-2 receptor.

### SARS-CoV-2 can infect liver cell lines in vitro

Previous studies have shown that ASGR1 is mainly expressed in liver and peripheral blood mononuclear cells [31–33]. Using the previously constructed SARS-CoV-2 pseudovirus system, we found that the pseudovirus could infect Huh-7 and HepG2 cells (Fig. 1F-1G).Therefore, we repeated the verification in Huh-7 and HepG2 cells. Both fluorescence microscopy (Fig. 3A) and flow cytometry (Fig. 3B) served to analyze GFP levels, while luciferase levels were assayed in parallel (Fig. 3C), proving that the SARS-CoV-2 pseudovirus could infect Huh-7 and HepG2 cells. Although results obtained by single-cell sequencing showed that ACE2 expression in human liver cells is low, there are reports that ACE2 can be stably expressed in commonly used *in vitro* liver cell lines. Therefore, we analyzed the expression of *ACE2* and *ASGR1* mRNAs and the corresponding proteins in Huh-7 and HepG2 cells by qPCR and western blotting, respectively (Fig. 3D-3E). The results showed that both Huh-7 and HepG2 cells expressed ACE2 and ASGR1, but the expression level of ASGR1 was significantly higher than that of ACE2. To further test whether the SARS-CoV-2 pseudovirus entered Huh-7 cells through ASGR1 or ACE2, we used CRISPR/Cas9 technology to knock out *ASGR1*, *ACE2*, or both. Next, we used SARS-CoV-2 pseudovirus to infect both cell lines. We found that in Huh-7 cells, neither knocking out *ACE2* nor *ASGR1* alone prevented the SARS-CoV-2 pseudovirus from infecting the cells, but when *ASGR1* and *ACE2* were knocked out simultaneously, it was evident that the efficiency of SARS-CoV-2 in infecting the cells was significantly reduced, which was confirmed by fluorescence microscopy, flow cytometry, and luciferase assays (Fig. 3F-H). This suggests that in Huh-7 cells, there may be two independent receptors mediating SARS-CoV-2 infections.

**Figure 3.**
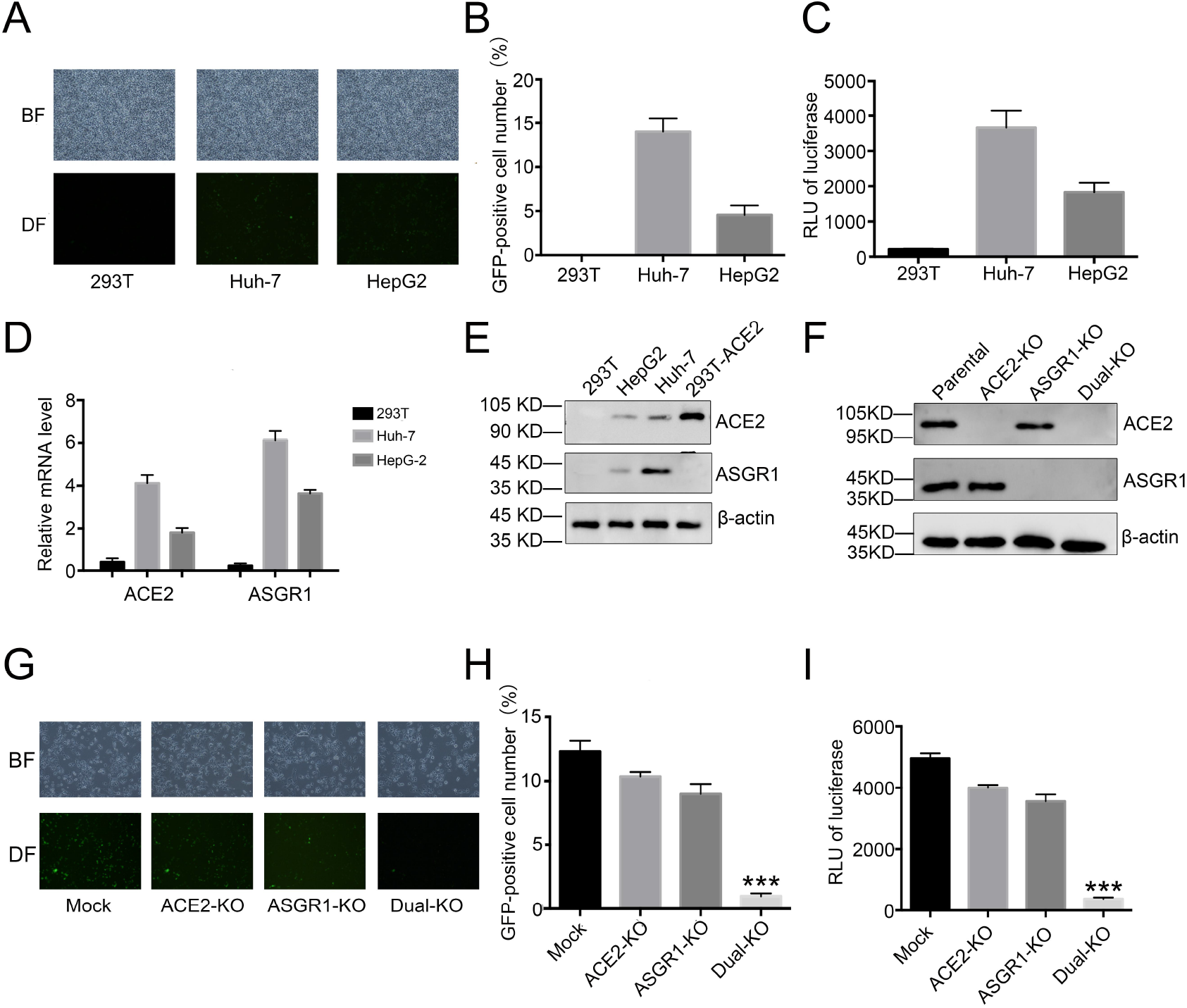
SARS-CoV-2 pseudovirus can infect liver cell lines through ASGR1 in *vitro*. (A-C) HEK293T, Huh-7 and HepG2 cells were infected with the GFP-labeled and luciferase SARS-CoV-2 virus pesudotype, and visualized by microcopy (A), the expression levels of GFP by cell flow cytometer (B), luciferase were detected by microplate (C) at 72 h post infection. (D, E) ACE2 and ASSGR1 expression were measured by qPCR (D) and western blot (E) in HEK293T, Huh-7 and HepG2 cells. (F) ACE2 and ASSGR1 expression were measured by western blot in Huh-7 which knock out *ASGR1*, *ACE2*, or both. (G-I) SARS-CoV-2 pseudovirus can infect Huh-7 cells through ACE2 or ASGR1. Huh-7 cells, which knock out *ASGR1*, *ACE2*, or both, were infected with the GFP-labeled and luciferase SARS-CoV-2 virus pesudotype, and visualized by microcopy (G), the expression levels of GFP by cell flow cytometer (H), luciferase were detected by microplate (I) at 72 h post infection. Each data represented the mean ± SD of three independent experiments (n=3) and were analyzed with T-test. ***, p < 0.001.

### SARS-CoV-2 infects immortalized liver cell lines and primary hepatocytes through ASGR1

Given that the expression level of ACE2 in normal liver tissue is much lower than 0.1% [14], while the expression level of ACE2 in liver tumor cell lines, such as HepG2 and Huh-7, is relatively high, the latter cannot fully reflect the condition of liver cells in the human body. Therefore, we considered further exploration of the immortalized liver cell line THLE-2 and primary liver parenchymal cells. We first determined the expression levels of ACE2 and ASGR1 proteins in THLE-2 and primary liver parenchymal cells; unlike Huh-7 and HepG-2 liver tumor cell lines, THLE-2 and primary liver parenchymal cells expressed nearly no ACE2, whereas ASGR1 was highly expressed (Fig. 4A-4B). This suggested that these two cell types may be a suitable cell system for the study of SARS-CoV-2 receptors in liver tissue. We first used SARS-CoV-2 pseudovirus to infect THLE-2 and primary liver parenchymal cells, and found that SARS-CoV-2 could infect both cell types (Fig. 4C-E).

**Figure 4.**
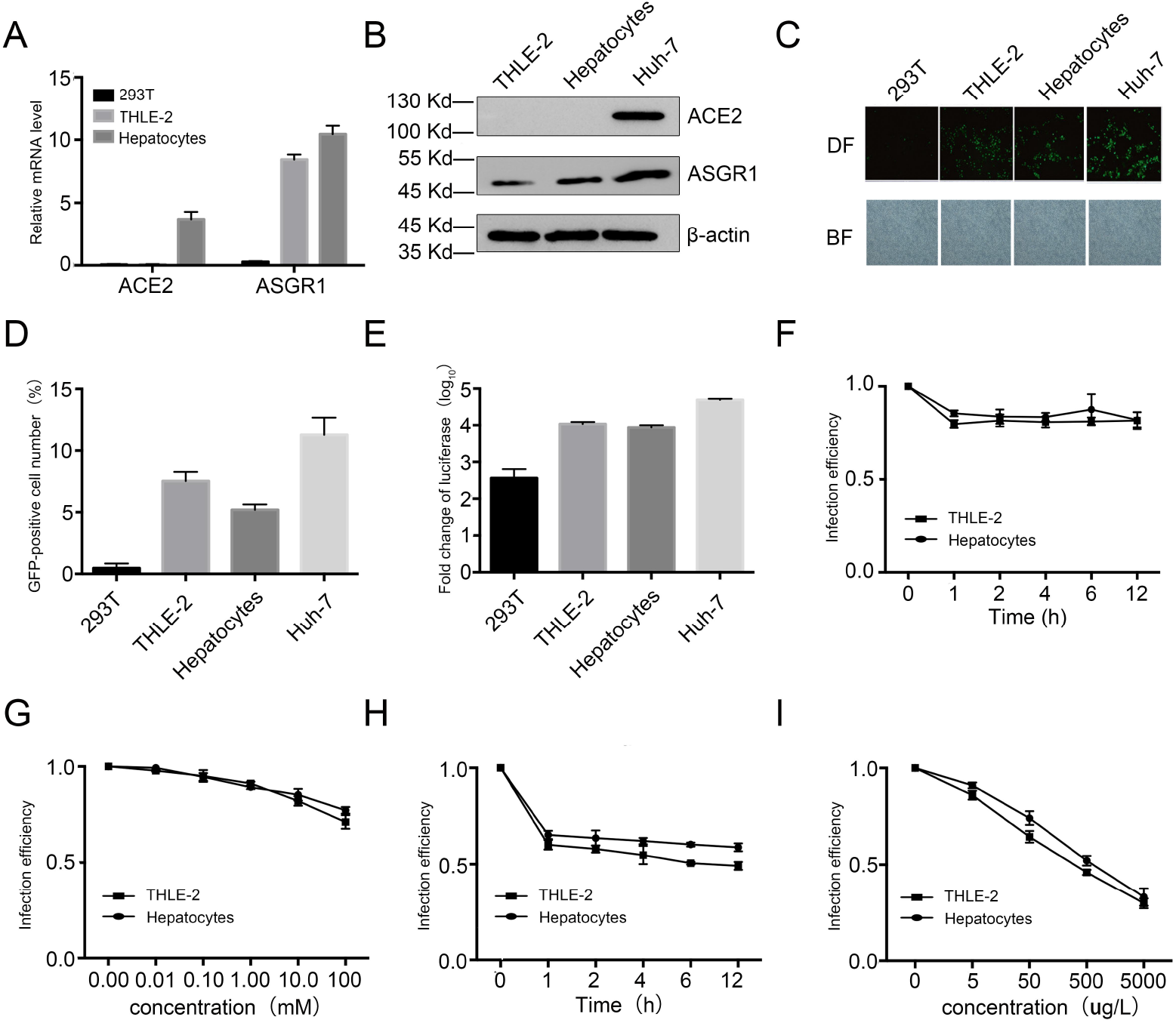
SARS-CoV-2 infects immortalized liver cell lines and primary hepatocytes through ASGR1. (A,B) ACE2 and ASGR1 expression were measured by qPCR (A) and western blot (B) in THLE-2 and primary hepatocytes cells. (C-E) SARS-CoV-2 pseudovirus can infect THLE-2 and primary hepatocytes cells. THLE-2 and primary hepatocytes cells were infected with the GFP-labeled and luciferase SARS-CoV-2 virus pesudotype, and visualized by microcopy (C), the expression levels of GFP by cell flow cytometer (D), luciferase were detected by microplate (E) at 72 h post infection. (F,G) Galactose cannot completely prevent the SARS-CoV-2 pseudovirus from infecting liver cells. The THLE-2 and primary hepatocytes cells were treated with 50 mM galactose for 0 h, 1 h, 2 h, 4 h, 8 h, 12 h (F) or with 0 nM, 10nM, 100nM, 1mM, 10mM, 100mM for 6h (G), then infected with SARS-CoV-2 virus pesudotype, and the infection efficiency was detected by luciferase. (H, I) ASGR1 monoclonal antibody can prevent SARS-CoV-2 pseudovirus from infecting liver cells. The THLE-2 and primary hepatocytes cells were treated with 500 ng/L anti-ASGR1 antibody for 0 h, 1 h, 2 h, 4 h, 8 h, 12 h (H) or with 0 ng/L 5 ng/L, 50 ng/L, 500 ng/L, 5 mg/L for 6h (I), then infected with SARS-CoV-2 virus pesudotype, and the infection efficiency was detected by luciferase. Each data represented the mean ± SD of three independent experiments (n=3) and were analyzed with T-test.

Given the short culture cycle of primary cells, it is not suitable to screen knockout cells through CRISPR-Cas9-dependent knockout. In THLE-2 and primary liver parenchymal cells, we chose to use galactose, which has been shown to interact with ASGR1 receptors, as well as ASGR1 monoclonal antibody to block ASGR1 binding, to determine whether SARS-CoV-2 pseudovirus infection could be prevented. Using this method, we found that in galactose-treated cells, the proportion of SARS-CoV-2-infected THLE-2 and primary liver parenchymal cells was only slightly decreased with increasing galactose concentrations, which may have been due to the combination of galactose and ASGR1; later, it was endocytosed into the cells and could not completely hinder the interaction between SARS-CoV-2 and ASGR1 (Fig. 4F-4G). At the same time, when ASGR1 monoclonal antibody was used to treat the cells, we found that the infection rate with SARS-CoV-2 was significantly decreased, which depended on the ASGR1 monoclonal antibody concentration used, but was independent of the incubation time (Fig. 4H-4I). These results indicate that ASGR1 is the main receptor for SARS-CoV-2 pseudovirus infection in immortalized and primary liver parenchymal cells, and that ASGR1 monoclonal antibody holds the potential to prevent SARS-CoV-2 infections.

### Soluble ASGR1 protein can prevent SARS-CoV-2 pseudovirus from infecting a various of cells

After confirming that ASGR1 is a potential new receptor for SARS-CoV-2, we tested whether it could be used for the prevention of SARS-CoV-2 infections. We first obtained soluble ASGR1 protein using 293T cells by transfection with an expression plasmid, followed by immunoprecipitation-based purification (data not shown). After that, we co-incubated with soluble ASGR1 protein and SARS-CoV-2 pseudovirus and then infected THLE-2 and primary liver parenchymal cells. As the concentration of soluble ASGR1 protein and time of co-incubation were increased, the efficiency of SARS-CoV-2 pseudovirus infection gradually decreased (Fig. 5A-5B). In addition to liver cells, which may depend on ASGR1 for SARS-CoV-2 entry, we wanted to further explore whether soluble ASGR1 could prevent susceptible cells featuring ACE2-mediated entry from being infected by SARS-CoV-2 pseudovirus. We repeated the experiment with H1299, 293T-ACE2, and Huh-7 cells, upon which we found that infection efficiencies in H1299 and Huh-7 cells were significantly decreased, while the decrease was not significant in the case of 293T-ACE2 cells (Fig. 5C-5D). The expression level of ACE2 in 293T-ACE2 cells was assumed to be much higher than under physiological conditions, thus potentially obscuring the treatment effects. Finally, because SARS-CoV-2 mainly infects epithelial cells of the respiratory tract, we decided to explore whether soluble ASGR1 protein had an effect on the proliferation and apoptosis of respiratory epithelial cells. The results showed that, regardless of the concentration, soluble ASGR1 had no significant effect on the proliferation and apoptosis of BEAS-2B cells (Fig. S1). This suggests that administration of soluble ASGR1 may be a safe and effective new anti-SARS-CoV-2 treatment approach.

**Figure 5.**
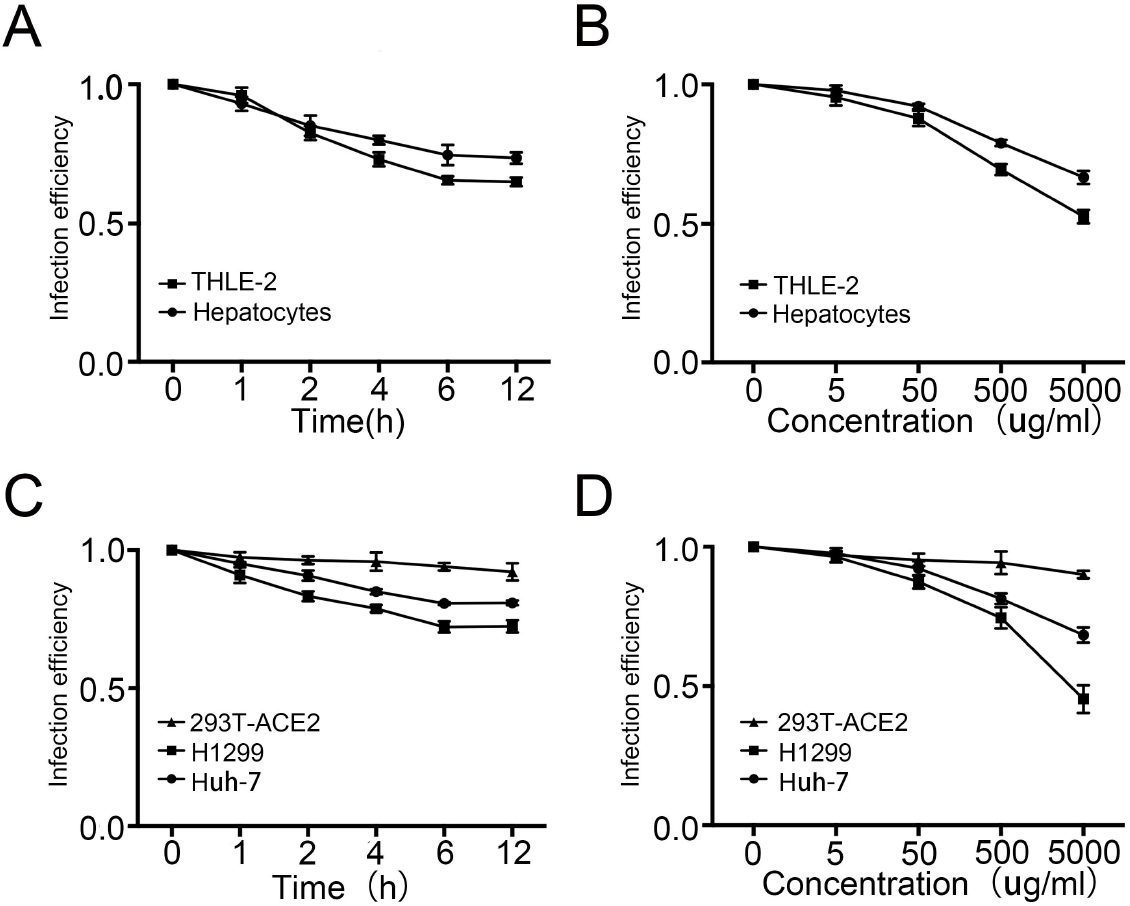
Soluble ASGR1 protein can prevent SARS-COV-2 pseudovirus from infecting a variety of cells. (A, B) soluble ASGR1 protein could prevent the SARS-CoV-2 pseudovirus from infecting liver cells. The THLE-2 and primary hepatocytes cells were infected with SARS-CoV-2 virus pesudotype which incubated with 500 ug/mL soluble ASGR1 for 0 h, 1 h, 2 h, 4 h, 8 h, 12 h (A) or with 0 ug/mL 5 ug/mL, 50 ug/mL, 500 ug/mL, 5000 ug/mL for 6h (B), and the infection efficiency was detected by luciferase. (C, D) soluble ASGR1 protein could prevent the SARS-CoV-2 pseudovirus from infecting cells which expressing ACE2. The Huh-7, H1299 and 293T-ACE2 cells were infected with SARS-CoV-2 virus pesudotype which incubated with 500 ug/mL soluble ASGR1 for 0 h, 1 h, 2 h, 4 h, 8 h, 12 h (C) or with 0 ug/mL, 5 ug/mL, 50 ug/mL, 500 ug/mL, 5000 ug/mL for 6h (D), then infected with SARS-CoV-2 virus pesudotype, and the infection efficiency was detected by luciferase. Each data represented the mean ± SD of three independent experiments (n=3) and were analyzed with T-test. *, p < 0.05;**, p < 0.01; ***, p < 0.001.

## Discussion

SARS-CoV-2 mainly infects cells of the respiratory system, but there is also increasing evidence that it can infect almost all major organs of the body, including the heart, brain, and liver [7–11]. Initially, researchers detected virus particles in epithelial cells of the respiratory tract, but recently, SARS-CoV-2 virus particles were found in pathological sections of various organs, such as the kidney, liver, and duodenum [7–11]. Clinical data shows that more than 50% of Covid-19 patients have severe liver damage and inflammation [8]. In addition, Covid-19 virus particles were also observed through electron microscopy in liver cells of Covid-19 patients’ autopsy [8]. This suggests to us that Covid-19 infection of liver cells may be the cause of severe liver damage in patients. However, the SARS-CoV-2 receptor ACE2, which was first discovered, is highly expressed in epithelial cells of the respiratory and digestive tracts. However, the expression level of ACE2 in liver cells is 0.1% [11,14]. Therefore, there may be non-ACE2 SARS-CoV-2 receptors in liver cells.

Although a number of studies on new receptors for the new coronavirus have been reported, the receptors found in most studies are mainly co-receptors for ACE2, such as host proteases transmembrane proteases serine 2 (TMPRSS2) [34] and Neuropilin-1 (NRP-1) [35]. The only known non-ACE2 receptors are AXL [11] and CD147 [36]. However, AXL and CD147 respectively mainly expresses lung cells and monocyte, which cannot explain the infection of other tissues by SARS-CoV-2. In some preprinted studies that have not been peer-reviewed, more information about the SARS-CoV-2 receptor is reported, but these receptor screenings are mainly concentrated in a few tool cells such as 293T, and there is no explanation about the physiological significance of the existence of non-ACE2 receptor of SARS-CoV-2 [37–39].

In this study, we conducted a gain-of-function screening experiment for the entry of SARS-CoV-2 into host cells, by which we uncovered ASGR1 as a new receptor, which was proven to be the main receptor for SARS-CoV-2 entry into liver cells. Previous studies confirmed that ASGR1 can promote HBV to infect human liver cells [40]. Moreover, due to its high expression in liver cells, ASGR1 is one of the current candidate targets for the treatment of liver cancer [41,42]. Conclusive evidence has been obtained by analyzing patient-derived liver samples from COVID-19 autopsies [7]. Compared with ACE2, ASGR1, which is highly expressed in liver cells, is more likely to be the main receptor for SARS-CoV-2 entry. In addition, we found that blocking ASGR1 with antibodies could significantly reduce SARS-CoV-2 infection. In addition, soluble ASGR1 protein could not only inhibit SARS-CoV-2 from infecting liver cells, but also prevented SARS-CoV-2 from infecting cells expressing ACE2 receptors, such as H1299 and 293T-ACE2 cells. We also found that soluble ASGR1 protein had no significant effect on the proliferation and viability of lung cells, suggesting that ASGR1 protein may be a potential candidate for the treatment of SARS-CoV-2.

In the present study, we could confirm that ASGR1 is the main receptor for SARS-CoV-2 entry into liver cells using various cell lines and primary cells. However, currently, we lack P3 laboratory conditions, and thus lack evidence of a correlation between authentic virus infection of liver cells and ASGR1. However, in the preprint study of bioRxiv, Tang et al. also showed that overexpression of ASGR1 in 293T could promote SARS-CoV-2 pseudovirus and authentic virus infections [37], suggesting that the latter may also enter cells through ASGR1 receptors.

Taken together, our findings demonstrate that ASGR1 plays an indispensable role in facilitating SARS-CoV-2 entry into liver cells. We anticipate that upon further validation and mechanistic elucidation, our findings will contribute to the development of therapeutic solutions for SARS-CoV-2 infection.

---

**Supplementary Figure 1.**
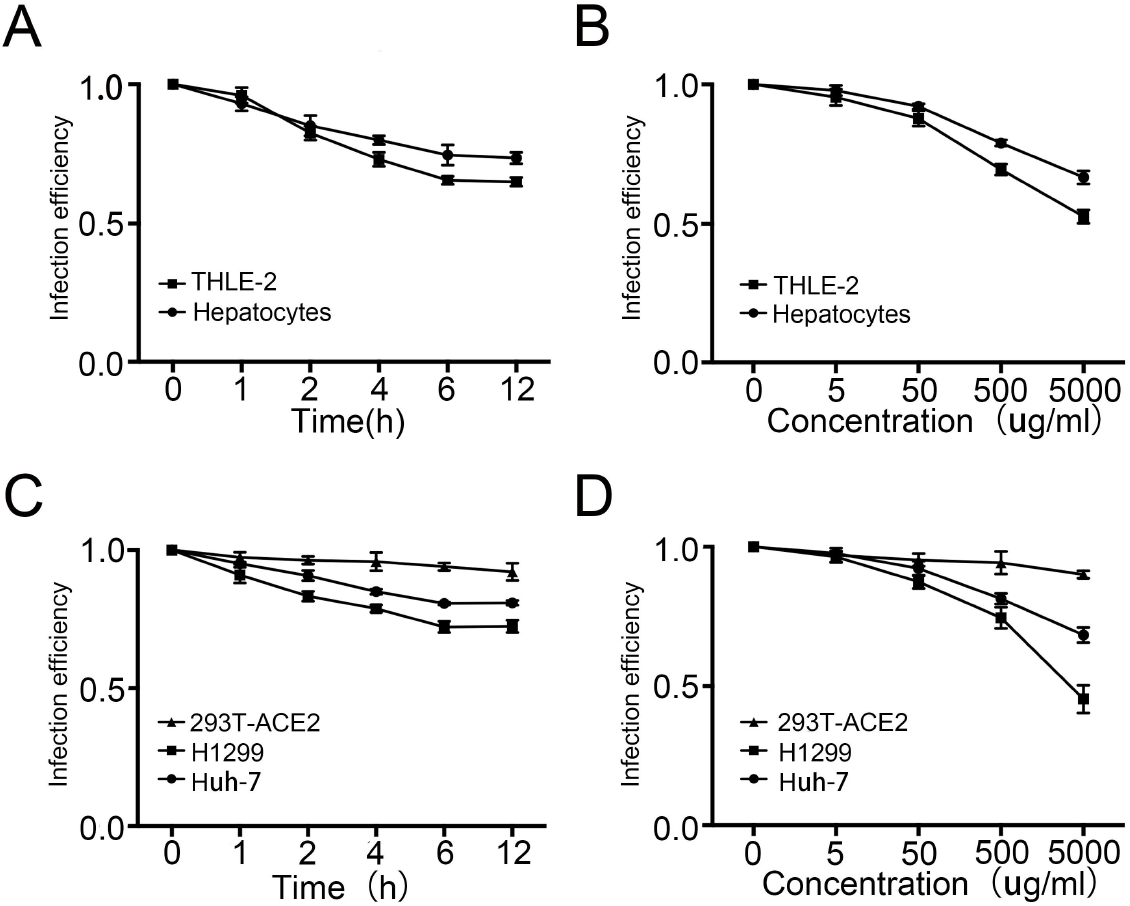
Effect of soluble ASGR1 protein on the proliferation and apoptosis of lung epithelial cells. (A) Cell proliferation of BEAS-2B cells incubated with 0 ug/mL, 5 ug/mL, 50 ug/mL, 500 ug/mL, 5000 ug/mL soluble ASGR1 protein for 24 h. cells was analyzed by CCK8. (B) Apoptosis of BEAS-2B cells incubated or not incubated with 500 ug/mL soluble ASGR1 protein for 24 h were measured by TUNEL staining followed by flow cytometry. Each data represented the mean ± SD of three independent experiments (n=3) and were analyzed with T-test.

